# Investigating the impact of paternal aging on murine sperm miRNA profiles and their potential link to autism spectrum disorder

**DOI:** 10.1101/2023.06.01.543326

**Authors:** Kazusa Miyahara, Misako Tatehana, Takako Kikkawa, Noriko Osumi

## Abstract

Paternal aging has consistently been linked to an increased risk of neurodevelopmental disorders, including autism spectrum disorder (ASD), in offspring. Recent evidence has highlighted the involvement of epigenetic factors. In this study, we aimed to investigate age-related alterations in microRNA (miRNA) profiles of mouse sperm and analyze target genes regulated by differentially expressed miRNAs (DEmiRNAs). Microarray analyses were conducted on sperm samples from mice at different ages: 3 months (3 M), over 12 M, and beyond 20 M. We identified 26 miRNAs with differential expression between the 3 M and 20 M mice, 34 miRNAs between the 12 M and 20 M mice, and 2 miRNAs between the 3 M and 12 M mice. The target genes regulated by these miRNAs were significantly associated with apoptosis/ferroptosis pathways and the nervous system. We revealed alterations in sperm miRNA profiles due to aging and suggest that the target genes regulated by these DEmiRNAs are associated with apoptosis and the nervous system, implying a potential link between paternal aging and an increased risk of neurodevelopmental disorders such as ASD. The observed age-related changes in sperm miRNA profiles have the potential to impact sperm quality and subsequently affect offspring development.

## Introduction

With the increasing trend of delayed parenthood, there has been a burgeoning focus on understanding the intricate relationship between aging and germline cells. While the effect of maternal aging on offspring, such as an elevated risk of miscarriage and Down syndrome, has garnered significant attention, the impact of male aging has received comparatively less recognition. However, recent research has revealed that advancing paternal age not only influences sperm morphology and motility but also has implications for DNA integrity and epigenetic factors^1–4^. These alterations in sperm composition due to paternal aging have been implicated in perturbations during the developmental process of offspring, potentially contributing to a spectrum of health issues^5^.

In recent decades, investigations into the origins of health and disease in future generations have gained paramount significance. The Developmental Origin of Health and Disease (DOHaD) concept has conventionally emphasized the impact of environmental factors on fetal development within the maternal uterus. However, there is now an emerging recognition that paternal factors also play a pivotal role in the risk of disease manifestation in children, giving rise to the concept of the Paternal Origin of Health and Disease (POHaD) concept^6,7^. For example, a study reported more than 800 differential DNA methylation regions between fathers who have children with autism spectrum disorder (ASD) and those who do not, suggesting that epigenetic alterations of sperm are key for ASD^8^. Paternal aging, in particular, has been linked to not only adverse incidents during pregnancy, such as miscarriage^9^ and stillbirth^10^ but also postnatal health issues, including low birth weight^11^, congenital heart defects^12^, and malignant tumors^13^. Although the concept of POHaD is relatively nascent, delving into this perspective holds the potential to provide insights into diseases that remain incompletely understood.

The concept of POHaD has emerged as a compelling framework, offering substantial evidence linking paternal factors to a range of neuropsychiatric disorders in offspring. Notably, advanced paternal age has been consistently demonstrated to increase the risk of schizophrenia^14^, bipolar disorder^15^, attention-deficit/hyperactivity disorder (ADHD)^16^, neurocognitive dysfunction^17^, and ASD^18^ in subsequent generations. The global surge in ASD^19^ has been influenced by a complex interplay of social and biological factors, encompassing updated diagnostic criteria, heightened social awareness, maternal drug exposure, and parental aging^18,20–22^. Remarkably, a meta-analysis of six million subjects across five countries revealed that paternal aging exerts a more pronounced impact on ASD risk than maternal aging^18^. Consequently, further exploration of the effects of POHaD may hold significant potential for providing invaluable insights into the pathophysiology of ASD.

The precise mechanism underlying the impact of paternal aging on the health of the next generation remains incompletely understood. *De novo* mutations, which are mutations that are not present in either parent but are observed in the child, have been associated with paternal aging and are believed to contribute to the increased disease risk in offspring. Specifically, *de novo* mutations of paternal origin have been found to accumulate with advancing paternal age, particularly in individuals diagnosed with ASD and schizophrenia^23,24^. Nevertheless, it is important to note that *de novo* mutations derived from aged fathers account for only a fraction, approximately 10-20%, of the overall risk of psychiatric disorders in children^25,26^. This suggests the involvement of additional factors that necessitate careful consideration in understanding the complete picture.

Epigenetic modifications in sperm have emerged as another plausible mechanism underlying the effects of paternal aging on offspring^27–31^. We have previously revealed age-related modifications in DNA methylation in mouse sperm^4^ and changes in histone modifications within the testis^32^, suggesting their potential influence on offspring development. MicroRNAs (miRNAs) are other crucial epigenetic factors that participate in gene expression regulation by inhibiting translation. These short noncoding RNA molecules (21-25 nucleotides) form complexes known as RNA-induced silencing complexes with their mRNA target sequences, ultimately suppressing translation^33^. It has been reported that aging affects the sperm profile of short noncoding RNAs, including miRNAs^34^, which might contribute to the aberrant sperm mRNA profile of fathers of ASD patients^35^. The biomedical research field actively investigates miRNAs due to their potential as biomarkers. Notably, aberrant miRNA profiles have been identified in individuals with ASD, with altered miRNA expression observed in various samples, such as the brain, blood, serum, and saliva^36–38^. Similar findings have been reported in animal models of ASD^39^. However, the precise mechanisms by which these changes in miRNA expression contribute to the pathogenesis of ASD remain unclear.

In a previous study on human sperm samples, age-related alterations in miRNA expression were reported, with certain miRNAs exhibiting increased levels and others showing decreased levels^2^. Additionally, alterations in mouse sperm miRNA expression, induced by factors such as diet and exercise, can impact the insulin sensitivity of female offspring^40^. Nevertheless, the comprehensive changes in overall miRNA profiles and the precise roles of the target genes regulated by these altered miRNAs remain elusive. Therefore, the primary objective of this study was to meticulously examine the expression profiles of miRNAs in sperm obtained from aged mice in comparison to their young counterparts. Furthermore, we aimed to unravel the potential functions of target genes of differentially expressed miRNAs (DEmiRNAs), shedding light on their roles in the context of paternal aging and potential implications for offspring mental health.

## Materials and methods

### Animals

C57BL/6J mice at 3 months of age were purchased from a breeder (Charles River Laboratories, Japan) and were raised to 12 or 20 months old at the Institute for Animal Experimentation at the Tohoku University Graduate School of Medicine; they were then used for miRNA and mRNA analysis. The mice were then sampled for microarray analyses at 3 months old (3 M), > 12 M or > 20 M (Fig. 1A). All animals were housed in standard cages in a temperature- and humidity-controlled room with a 12-hour light/dark cycle (lights on at 8 am), and they had free access to standard food and water. All experimental procedures were approved by the Ethics Committee for Animal Experiments of the Tohoku University Graduate School of Medicine (#2017-MED210, #2019-MED280-03, 2020BeLMO-005-01 and 2020BeA-015-01) and complied with the ARRIVE guidelines. The animals were treated according to the National Institutes of Health guidance for the care and use of laboratory animals.

**Figure 1.**
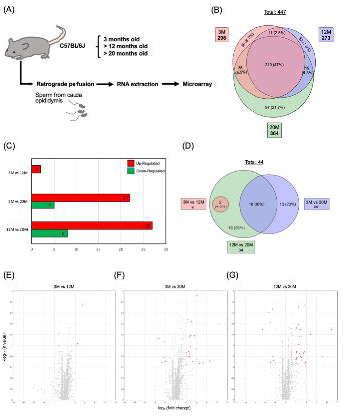
Analyses of DEmiRNAs in sperm samples of different ages. **(A)** Experimental design for analyzing sperm miRNAs. Sperm samples were collected from the cauda epididymides of three C57BL/6J mice at 3 M, > 12 M, and > 20 M via retrograde perfusion. Further details can be found in the text. (**B**) Venn diagram showing the number of miRNAs expressed in sperm samples from mice aged 3 M, 12 M and 20 M. A total of 447 miRNAs were expressed. Black letters indicate the numbers of common miRNAs and their percentages against the total numbers. (**C**) DEmiRNAs were defined as those showing > 2-fold up- or downregulation and *P* < 0.05. Red bars indicate upregulated miRNAs, and green bars indicate downregulated miRNAs. (**D**) Venn diagram showing DEmiRNAs in sperm samples in the 3 M vs. 12 M, 3 M vs. 20 M, and 12 M vs. 20 M comparisons. A total of 46 DEmiRNAs were identified. Black letters indicate the numbers of common miRNAs and their percentages against the total numbers. (**E-G**) Volcano plots showing DEmiRNAs in sperm samples from mice aged 3 M vs. 12 M (**E**), 3 M vs. 20 M (**F**), and 12 M vs. 20 M (**G**). DEmiRNAs were defined as those showing > 2-fold up- or downregulation and *P* < 0.05. Red dots indicate upregulated miRNAs, and green dots indicate downregulated miRNAs.

### Sperm collection for miRNA and mRNA analyses

For microarray analyses, three mice of each age were used. Because of visible age-related changes, the experimenter could not blind the animals to their age. Mice were anesthetized with isoflurane (MSD Animal Health) and perfused with PBS (pH 7.4) to remove blood. After perfusion, the cauda epididymis was removed from fat and other connective tissues to collect sperm by retrograde perfusion^41–43^. Sperm were centrifuged on a 27% Percoll solution (2005-001 or 2001-017, NK system). The pellet was resuspended in PBS and recentrifuged (400×g for 2 min) to remove excess Percoll. The purity of the pellet was confirmed by microscopy. The samples were then subjected to miRNA and mRNA extraction.

### RNA extraction and microarray analysis

Sperm samples were subjected to extraction of total RNA, including miRNA or mRNA, by using a miRNeasy Mini Kit (217004, Qiagen). We confirmed whether total RNA, including miRNA, was sufficiently collected with an Agilent Small RNA Reagent Kit (5067-1549) and a 2100 Bioanalyzer Instrument (G2939BA). RNA samples were subjected to microarray analyses. A Clariom™ D Assay for mice (902513, Thermo Fisher Scientific) and a GeneChip™ miRNA 4.0 Array (902412, Thermo Fisher Scientific) were used for mRNA and miRNA analyses, respectively. Transcript expression values were normalized using the Signal Space Transformation (SST-RMA) method. Raw intensity data (i.e., CEL files) were analyzed by using Transcriptome Analysis Console (TAC) Software version 4.0.2 (Affymetrix Transcriptome Analysis Console Software, RRID: SCR_018718). We identified DEmiRNAs based on > 2-fold up- or downregulation along with a *P* value < 0.05. The microarray data have been deposited into the Gene Expression Omnibus database under accession number GSE232620 and are available at the following URL: https://www.ncbi.nlm.nih.gov/geo/query/acc.cgi?acc=GSE232620. Pathway analyses were conducted using the Reactome pathway analysis package in R^44^. Networks were constructed using Cytoscape^45^, which visualized relationships between enriched pathways and genes. Validated miRNA target genes were obtained from TAC Software. Pearson’s correlation test was utilized to analyze the correlation between miRNAs (i.e., *mmu-let-7b-5p*, *mmu-miR-10a-5p* and *mmu-miR-24-3p*) and their target genes related to apoptosis (i.e., *Ppp3r1* and *Bcl2l11*). We also compared the expression levels of the five miRNAs that have previously been reported as candidate miRNAs (i.e., miR-10a-5p, miR-146a-5p, miR-34c-5p, miR-143-3p, and miR-146b-5p) transmitted from sperm to fertilized eggs^46^. In these analyses, *P* < 0.05 was considered to indicate significance. We used Bonferroni’s multiple comparison test to verify the significance of the microarray results.

### Analyses exploring the possible functions of target genes of altered miRNAs

To investigate the possible functions of the target genes of DEmiRNAs in relation to ASD, we utilized the autism-related gene database SFARI^47^. We constructed a network of miRNA-target gene interactions by using Cytoscape. Additionally, we conducted Pearson’s correlation test between the expression levels of miRNAs (*mmu-miR-466j*, *mmu-miR-24-3p*, *mmu-miR-690*, and *mmu-let-7b-5p*) and their target SFARI genes (*Oxtr*, *Gabrb2*, *Ctnnb1*, and *Grik2*).

## Results

### Age-related changes in the expression of miRNAs in mouse sperm

We first analyzed age-related changes in sperm miRNA expression patterns. A total of 298, 273, and 364 miRNAs were expressed in sperm samples of the 3 M, 12 M, and 20 M groups, respectively (Fig. 1B). There were 221 miRNAs commonly expressed in the 3 M and 12 M groups and 248 miRNAs commonly expressed in the 3 M and 20 M groups. In the 12 M and 20 M groups, 229 miRNAs were commonly expressed, and 210 miRNAs were expressed in all three groups (Fig. 1B).

The miRNAs were then compared between the two groups. We considered the miRNAs that showed > 2-fold up- or downregulation along with *P* < 0.05 in expression as significantly differentially expressed. Two miRNAs were upregulated in the sperm samples collected from the 12 M group compared with the sperm samples from the 3 M group, 21 miRNAs were upregulated and 5 miRNAs were downregulated in the 3 M samples vs. the 20 M samples, and 26 and 8 miRNAs were up- and downregulated, respectively, in the 12 M vs. 20 M samples (Fig. 1C-G). The DEmiRNAs in each comparison are summarized in Supplementary Table S1.

No overlap was observed between the miRNAs that exhibited alterations in the 3 M vs. 12 M comparison and those in the 3 M vs. 20 M comparison (Fig. 1D). In the 12 M vs. 20 M comparison, two miRNAs that displayed changes in the 3 M vs. 12 M comparison were also found to be altered (4.5% of the total altered miRNAs). Furthermore, out of the 26 miRNAs that showed alterations in the 3 M vs. 20 M comparison, 16 were similarly altered in the 12 M vs. 20 M comparison (36% of the total altered miRNAs). Notably, this subset of miRNAs included *let-7b-5p*, *miR-10a-5p*, and *miR-24-3p,* which target apoptosis-related genes (Fig. 1D; Supplementary Table S2). These results indicated a distinct age-associated shift in the sperm miRNA profiles, particularly in the 3 M vs. 20 M and 12 M vs. 20 M age group comparisons.

### Targets of age-related miRNAs in mouse sperm

Next, the miRNAs significantly altered in the comparisons between the groups were listed, and their target genes (as listed in Supplementary Table S2) were subjected to enrichment analyses by using an R package for Reactome Pathway Analysis^44^ to identify biological pathways related to the DEmiRNAs. Although the 3 M vs. 12 M comparison did not have any pathways that were significantly different (Fig. 2A), the comparison between the 3 M and 20 M samples yielded differential pathways including “Dimerization of procaspase-8”, “CASP8 activity is inhibited”, “Regulation by c-FLIP”, “Apoptosis”, “Caspase activation via Death Receptors in the presence of ligand”, and “Caspase activation via the extrinsic apoptotic signaling pathway (Fig. 2B). Most of these pathways were related to the regulation of apoptosis.

**Figure 2.**
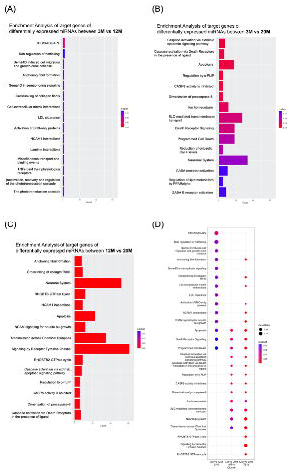
Pathway analyses of miRNA target genes. Pathway analyses were performed on miRNA target genes that significantly altered the sperm samples of 3 M vs. 12 M, 3 M vs. 20 M, and 12 M vs. 20 M using the Reactome pathway analysis package in R. The number of genes associated with each pathway is shown on the horizontal axis in (**A-C**). The gene ratio in (**D**) indicates the percentage of genes associated with each pathway among the genes listed as targets of the miRNAs.

In our comparison between 12 M and 20 M samples, we also found several pathways that showed significant differences. These included the same apoptosis-related pathways, such as “Caspase activation via Death Receptors”, “Dimerization of procaspase-8” and “Apoptosis”, as well as other pathways such as “Signaling by Receptor Tyrosine Kinases”, “Transmission across Chemical Synapses”, “NCAM signaling for neurite out-growth” and “Neuronal System” (Fig. 2C). Although those pathways related to the nervous system were not significantly enriched in the comparison between 3 M vs. 20 M samples (Fig. 2B), other significant pathways in the comparison between 3 M vs. 20 M samples overlapped with those that were also significantly altered between 12 M and 20 M samples (Fig. 2D). Furthermore, four genes related to ferroptosis, another type of cell death, *Acsl4*, *Hmox1*, ß*Slc11a2*, and *Slc3a2,* were identified as target genes of altered miRNAs between 3 M vs. 20 M samples. Among them, *Acsl4* and *Hmox1* were also target genes in the comparison between 12 M and 20 M samples. Therefore, our findings suggest that aging may be linked with cell death during spermatogenesis. In addition, those impacts on unique pathways seem to be caused by many miRNAs, not by one or two specific miRNAs. For example, 9 and 13 DEmiRNAs that regulate apoptosis-related genes were identified in the 3 M vs. 20 M and 12 M vs. 20 M comparisons, respectively (Supplementary Tables S3 and S4).

### Changes in the mRNA expression of age-associated miRNA target genes

We measured the expression levels of target genes using a commercial microarray kit, and the top 10 enriched pathways and genes were visualized using Cytoscape^45^ (Fig. 3). Among these pathways, we focused on apoptosis-related pathways (such as “Caspase activation via Death Receptors”, “Dimerization of procaspase-8” and “Apoptosis”) and investigated whether the expression levels of the target genes regulated by altered miRNAs were affected by aging (Supplementary Table S5). The network representations in Fig. 3 revealed that the expression levels of most genes, including *Casp9*, *Tnfsf10*, and *Fadd*, exhibited minimal changes. However, the expression levels of *Ppp3r1,* regulated by *mmu-let-7b-5p,* and *Bcl2l11,* regulated by *mmu-miR-10a-5p* and *mmu-miR-24-3p,* increased with sperm aging. Notably, *Ppp3r1* showed a significant increase in 20 M mice compared to 3 M mice (fold change = 1.5, *P* = 0.029). To validate the association between DEmiRNAs and changes in the gene expression levels of regulated genes, we conducted Pearson’s correlation tests. The results demonstrated a significant positive correlation between the expression levels of *Ppp3r1* and *mmu-let-7b-5p* (Supplementary Fig. S1A: *R* = 0.840, *P* = 0.005). Although *Bcl2l11* also exhibited a positive trend with *mmu-miR-10a-5p* and *mmu-miR-24-3p,* the correlation did not reach statistical significance (Supplementary Fig. S1B and S1C; *mmu-miR-10a-5p*: *R* = 0.646, *P* = 0.060; *mmu-miR-24-3p*; *R* = 0.613, *P* = 0.079).

**Figure 3.**
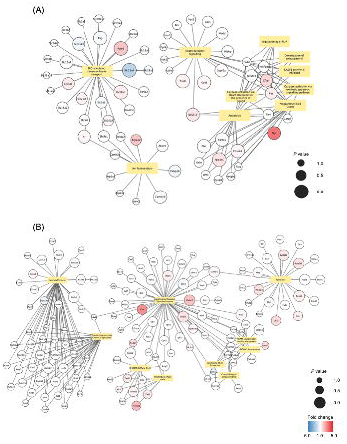
Network of biological pathways and genes. Network of significantly enriched pathways and target genes belonging to those pathways when comparing 3 M vs. 20 M (**A**) and 12 M vs. 20 M (**B**). The node color indicates the fold change in gene expression. The node size indicates the significance of the differential expression; larger sizes indicate smaller *P* values.

### ASD-related target genes of altered miRNAs

Although paternal aging is a risk factor for ASD^18,22^, the impact of DEmiRNAs with aging on ASD-related genes remains unknown. Therefore, we focused on genes associated with ASD and analyzed whether the target genes of the age-related miRNAs might have some relation with ASD. We searched the autism-related gene database SFARI^47^ and identified an overlap of 28 and 38 SFARI genes with the target genes of DEmiRNAs in 3 M vs. 20 M samples and 12 M vs. 20 M samples, respectively (Supplementary Table S6). No SFARI genes were identified as miRNA targets in comparisons with 3 M and 12 M samples. The relationship between miRNAs and SFARI genes is visualized in Supplementary Fig. S2 as a network. We focused on four SFARI genes whose expression levels based on our microarray analysis differed significantly between the groups (i.e., *Oxtr*, *Gabrb2*, *Ctnnb1*, and *Grik2*) and confirmed correlations of their expression levels with those of their regulatory miRNAs. These included significant negative correlations between *mmu-miR-466j* and *Oxtr* (Fig. 4A; *R* = -0.701, *P*= 0.035) and between *mmu-miR-24-3p* and *Gabrb2* (Fig. 4B; *R* = -0.707, *p*=0.033), while there was a tendency of positive correlation between *mmu-miR-690* and *Ctnnb1* (Fig. 4C; *R* = 0.615, *P*=0.078). The negative correlation between *mmu-let-7b-5p* and *Grik2* was not significant (Fig. 4D; *R* = -0.246, *P* = 0.523).

**Figure 4.**
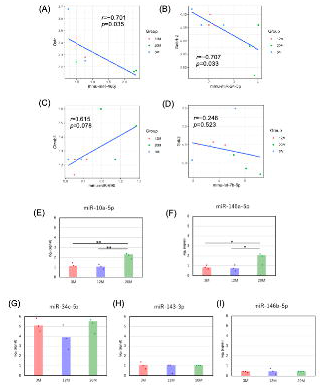
Expression levels of specific miRNAs. (**A-D**) Correlation analyses between SFARI genes whose expression levels varied between groups and their regulatory miRNAs. (**A**) *mmu-miR-466j* vs. *Oxtr*, (**B**) *mmu-miR-24-3p* vs. *Gabrb2*, (**C**) *mmu-miR-690* vs. *Ctnnb1*, (**D**) *mmu-let-7b-5p* vs. *Grik2*. (**E-I**) Expression scores in this microarray analysis of miRNAs that were previously reported to be inherited from sperm to fertilized eggs^46^. (**E**) *miR-10a-5p*, (**F**) *miR-146a-5p*, (**G**) *miR-34c-5p*, (**H**) *miR-143-3p*, and (**I**) *miR-146b-5p*. The bars show the mean of the signals of the three samples in each group. Bonferroni’s multiple comparison test was used to test for significant differences. \**P* < 0.05, ** *P* < 0.01.

### Changes in the expression of miRNAs that may be transmitted from sperm to fertilized eggs

Since some miRNAs altered by aging could be transferred to the fertilized egg and affect its quality, we finally focused on candidate miRNAs that can be carried from sperm to the fertilized egg based on previous mouse research comparing the miRNA profiles of sperm, oocytes and fertilized eggs^46^. We compared the expression levels of the miRNAs *miR-10a-5p*, *miR-146a-5p*, *miR-34c-5p*, *miR-143-3p*, and *miR-146b-5p* among the 3 M, 12 M, and 20 M groups based on our microarray data. Our results revealed significant upregulation of *miR-10a-5p* and *miR-146a-5p* in the 20 M mice (Fig. 4E-I). This finding highlights the potential significance of *miR-10a-5p* and *miR-146a-5p* as candidate sperm miRNAs transmitted to the zygote and the possible implications for subsequent offspring development.

## Discussion

### Changes in the expression of miRNAs with aging: Implications for cell death or the nervous system

Our analyses of miRNAs in sperm samples obtained from mice at different ages (3 M, 12 M, and 20 M) revealed distinct patterns of miRNA expression. Notably, when comparing the miRNA profiles between the youngest (3 M) and oldest (20 M) groups, we observed significant associations with pathways related to apoptosis (Fig. 2B). This finding aligns with previous studies demonstrating an increase in apoptosis and DNA damage in human and mouse sperm with advanced age^48,49^.

Through our analyses, we identified miRNA target genes that both promote (e.g., *Bcl2l11*, *Casp9*, and *Ppp3r1*) and inhibit (e.g., *Tnfsf10*, *Fadd*, and *Fas*) apoptosis. The expression levels of these genes play critical roles in regulating apoptosis. For instance, decreased expression of *Bcl2l11* and *Casp9* has been associated with suppression of sperm apoptosis^50^. Furthermore, exposure to recombinant tumor necrosis factor-related apoptosis-inducing ligand (encoded by *Tnfsf10*) can modulate sperm apoptosis^51^. Our findings revealed variations in the expression levels of certain genes, including *Bcl2l11* and *Ppp3r1,* across different age groups, suggesting that changes in the miRNA profile may contribute to the aberrant expression of these target genes.

Given that 20 M represents a highly advanced age, it is important to consider that the aging process is accompanied by a multitude of factors, including increased oxidative stress, lipid peroxidation, iron accumulation, and chronic inflammation. These factors may collectively contribute to heightened aging-related stress, which may not only promote apoptosis and ferroptosis but also increase the occurrence of necrosis. Our analysis revealed the presence of pathways related to apoptosis and ferroptosis in the comparison between the 12 M and 20 M groups (Fig. 2C), further supporting the notion that changes in the miRNA profiles of sperm resulting from paternal aging can influence different forms of cell death.

Our investigation uncovered several intriguing aging-related pathways, such as “Signaling by Receptor Tyrosine Kinases”, “Transmission across Chemical Synapses”, “Neuronal System” and “NCAM signaling for neurite out-growth”. The consistency of these findings is particularly noteworthy, considering the shared expression of many genes in both the brain and testes^52^. Examples of such shared genes are those related to calcium signaling, a crucial process in both neurons and sperm. Calcium channels play a central role in neurotransmission in the brain^53^ and are essential for sperm motility and capacitation^54^. In our study, we identified target genes involved in calcium signaling, including *Camkk1* and *Cacna2d2*. *Cacna2d2* encodes the alpha-2/delta subunit of the voltage-dependent calcium channel complex, while *Camkk1* encodes Ca^2+^calmodulin-dependent protein kinase α, both of which are essential for proper calcium signaling in sperm^55,56^. Consequently, aberrant expression of these genes due to miRNAs may disrupt sperm motility and capacitation^54^, both of which are also associated with paternal aging^1,57^. Thus, our findings suggest that changes in signaling pathways related to calcium signaling and other pathways identified in sperm from aged mice could contribute to the observed decline in sperm quality in aged males.

### Changes in the expression levels of genes caused by altered miRNAs with aging

Our study provides compelling evidence that altered miRNAs impact the expression levels of specific genes in mouse sperm. Particularly noteworthy were the age-related changes in the expression levels of the two apoptosis-related genes *Ppp3r1* and *Bcl2l11*. These changes demonstrated proportional associations with the expression levels of their regulatory miRNAs, namely, *mmu-let-7b-5p*, *mmu-miR-10a-5p*, and *mmu-miR-24-3p*, as quantitatively determined by microarray analyses. Previous research has indicated that upregulation of these three miRNAs is associated with apoptosis^58–60^. For instance, Huang, Y. et al. demonstrated that *let-7b-5p* promotes apoptosis and that knockdown of *let-7b-5p* suppresses apoptosis in cultured human cells^58^. Moreover, *mmu-miR-10a-5p* has been suggested to be transmitted to a fertilized egg^46^, which may cause additional effects on the transcriptome after fertilization. These findings lend further support to our hypothesis that these three miRNAs may indeed contribute to sperm apoptosis.

The positive correlations observed between the regulatory miRNAs and their target genes (Supplementary Fig. S1A-C) suggest the possibility of upregulation of *Ppp3r1* and *Bcl2l11* by these miRNAs. While miRNAs are commonly known for their roles in repressing gene expression, in some cases, there are negative correlations. For example, in cells at G0 phase, such as oocytes, miRNAs can promote gene expression^61^. Although specific studies investigating the effects of these miRNAs on mRNA upregulation in sperm are currently limited, it is plausible that the target genes influenced by these miRNAs could undergo upregulation in sperm obtained from aged mice, potentially leading to the induction of apoptosis.

### Paternal aging and ASD risk of offspring

We further investigated the impact of altered miRNAs on autism-related SFARI genes through microarray analyses. Although changes in the expression levels of SFARI genes were generally limited, intriguing associations were found between the expression levels of *Oxtr* and *Gabrb2* and their regulatory miRNAs. *Oxtr* encodes an oxytocin receptor, a crucial component in the regulation of social behavior^62^. A meta-analysis previously identified four polymorphisms in the human *OXTR* gene (rs7632287, rs237887, rs2268491, and rs2254298) that are linked to ASD^63^. Similarly, *Gabrb22* encodes the gamma-aminobutyric acid (GABA) type A receptor subunit beta2, an important component of the inhibitory system in the brain^64^. Disruptions of the GABA system have been implicated in ASD risk^65^. Polymorphisms in *GABRB2*, such as rs2617503 and rs12187676, have been implicated in ASD risk^66^. Although the roles of *Oxtr* and *Gabrb2* in germline cells are not well understood, their crucial functions in brain development are unknown. Aberrant expression of these genes due to miRNA changes in sperm may provide partial insights into the relationship between paternal aging and the increased risk of ASD in offspring.

## Limitations

While our study provides valuable insights into the role of miRNAs in paternal aging and their potential association with ASD risk, several limitations should be acknowledged. First, to fully understand the physiological functions of these miRNAs in fertilized eggs and their relationship to ASD, additional investigations involving techniques such as miRNA injection or knockdown in fertilized mouse eggs are necessary. Despite the relatively low abundance of miRNAs in sperm compared to fertilized eggs, they still hold promise as potential biomarkers for predicting the physical conditions of offspring. In addition, it is important to acknowledge that our study focused on miRNAs and their target genes, but other regulatory mechanisms and factors may also contribute to the development of ASD. Exploring additional epigenetic modifications and factors that influence gene expression could provide a more comprehensive understanding of the paternal contribution to ASD.

## Conclusion

In this study, we aimed to investigate epigenetic changes in male germline cells induced by aging and their potential impact on the development of the next generation. Our findings shed light on the alterations in the miRNA profile of sperm as a result of aging, which can subsequently lead to changes in gene expression levels. In addition to the well-established importance of the maternal environment in the DOHaD concept, our study highlights the growing relevance of investigating the effects of POHaD. By exploring both maternal and paternal contributions, we can gain comprehensive insights into the complex interplay among genetics, epigenetics, and environmental factors in shaping the health outcomes of future generations.

## Data availability statement

Microarray data were deposited into the Gene Expression Omnibus database under accession number GSE232620 and are available at the following URL: https://www.ncbi.nlm.nih.gov/geo/query/acc.cgi?acc=GSE232620.

## Acknowledgments

We would like to express our sincere gratitude to Prof. Eileen McLaughlin at the University of Wollongong for the online discussions and many helpful suggestions. We would also like to express our deep gratitude to Prof. Takashi Shinohara at Kyoto University Graduate School of Medicine and Dr. Hiroko Hamada at Tohoku University Graduate School of Medicine for their various advice on experiments and teaching experimental skills. We would also like to express our deepest gratitude to Prof. Yasuhisa Matsui, Prof. Takahiro Arima, and Mr. Jasper Germeraad at Tohoku University Graduate School of Medicine for their critical reading and valuable feedback, as well as our excellent technical staff member, Ms. Sayaka Makino, for taking care of the experimental mice and preparing the experimental environment.

## Authors’ contributions

MT and NO designed the study. MT, KM, and TK performed the experiments. MT and KM undertook the bioinformatic and statistical analyses. KM, MT and NO wrote the manuscript. All authors contributed and approved the final manuscript.

