## Supplementary Figures for "Investigating the impact of paternal aging on murine sperm miRNA profiles and their potential link to autism spectrum disorder"

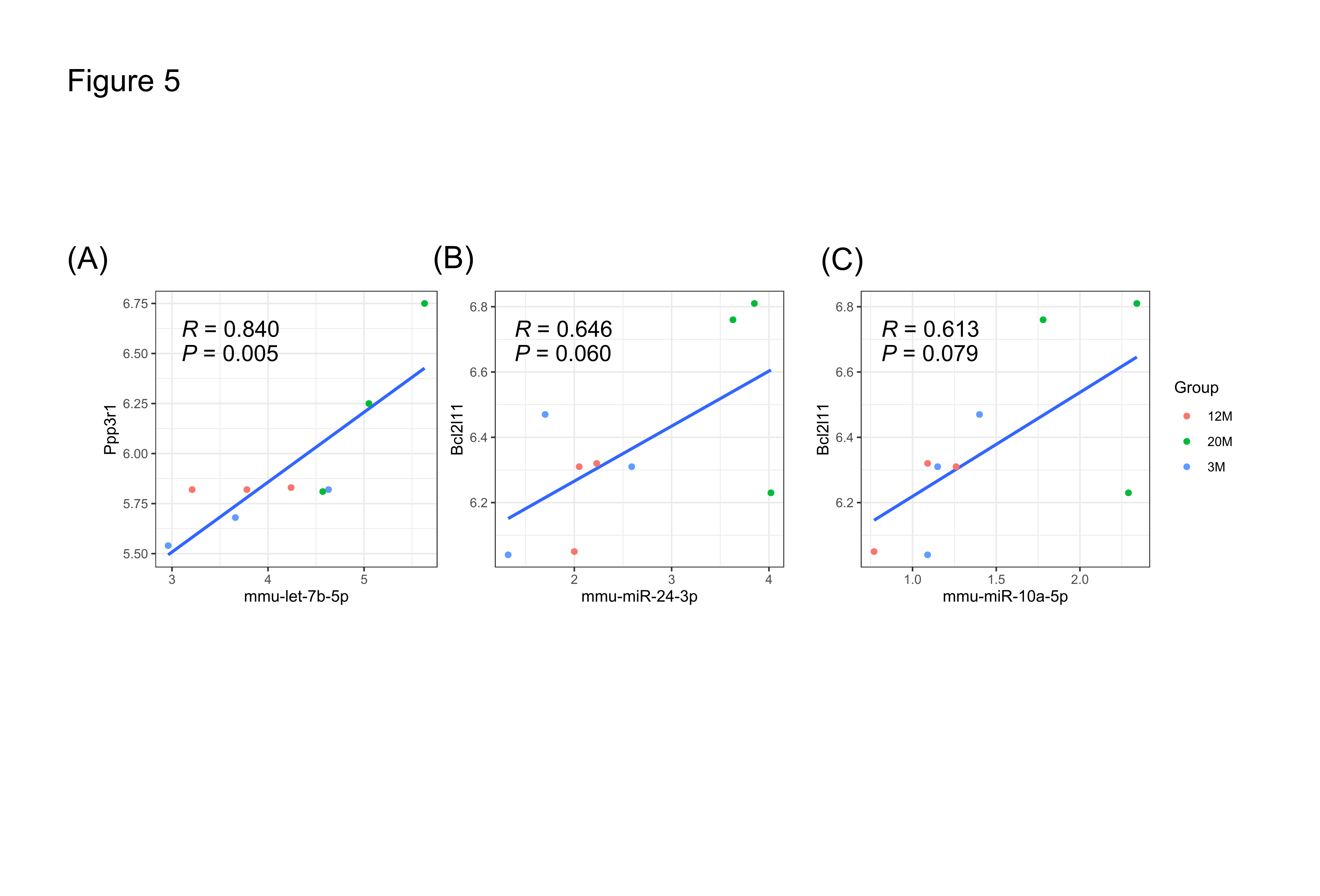
**Supplementary Figure S1**

**Supplementary Figure S1.** Correlation analysis between apoptosis-related genes, *Ppp3r1* and *Bcl2l11*, and their regulator miRNAs. Scatter plot shows the correlation between *Ppp3r1* and *mmu-let-7b-5p*　(**A**), *Bcl2l11* and *mmu-miR-10a-5p* (**B**), and *Bcl2l11* and *mmu-miR-24-3p* (**C**).

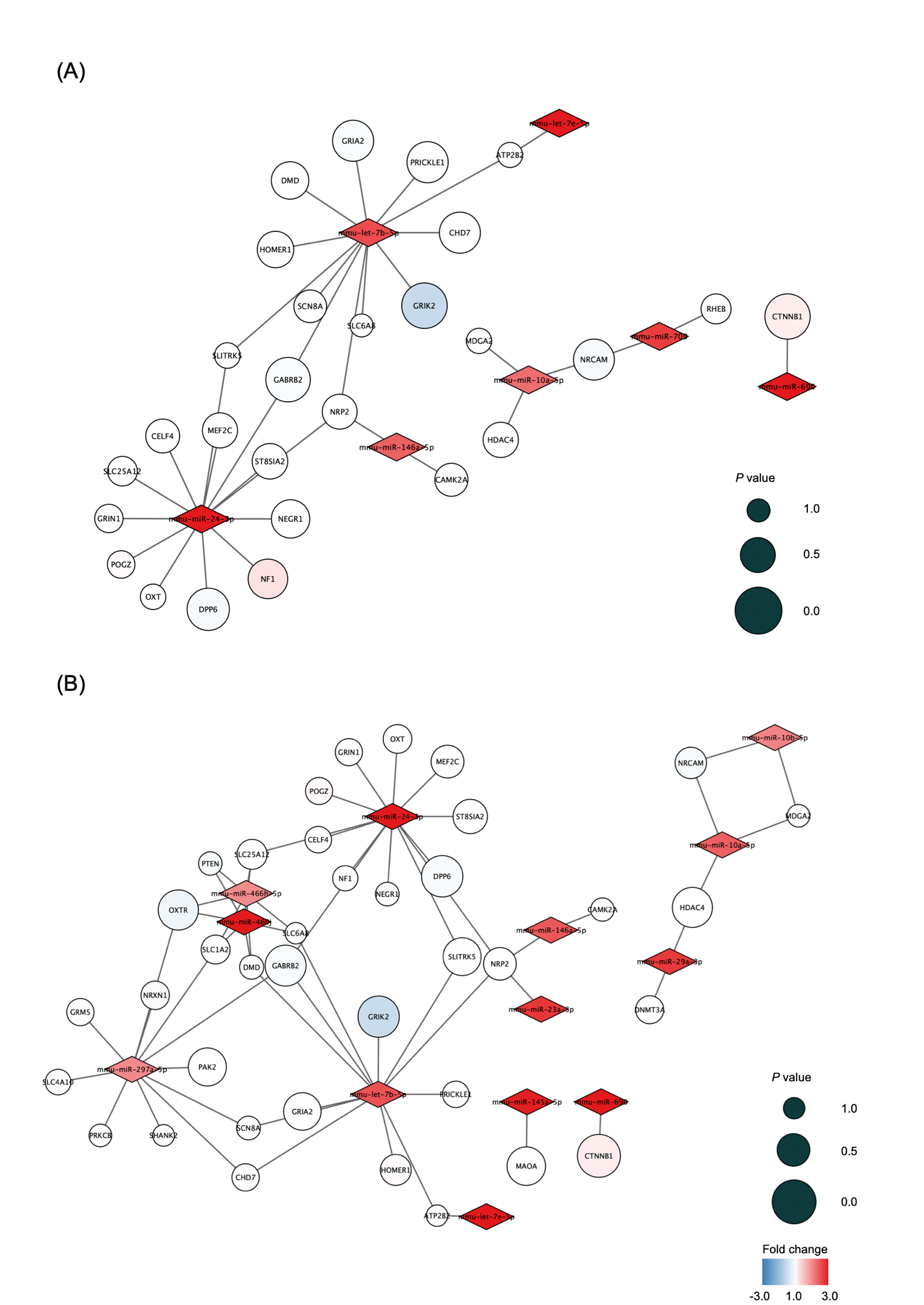
**Supplementary Figure S2**

**Supplementary Figure S2.** Network of miRNAs and SFARI genes. Network of significantly altered miRNAs and their target autism-related SFARI genes when comparing 3M vs 20M (**A**) and 12M vs 20M (**B**). Node color indicates the fold change of the miRNA and target gene expression. Node size indicates the significance of the differential expression; larger sizes indicate smaller *P* values.
